# Integrative analyses of splicing in the aging brain: role in susceptibility to Alzheimer’s Disease

**DOI:** 10.1101/174565

**Authors:** Towfique Raj, Yang I. Li, Garrett Wong, Satesh Ramdhani, Ying-chih Wang, Bernard Ng, Minghui Wang, Ishaan Gupta, Vahram Haroutunian, Eric E. Schadt, Bin Zhang, Tracy Young-Pearse, Sara Mostafavi, Pamela Sklar, David Bennett, Philip L. De Jager

## Abstract

We use deep sequencing to identify sources of variation in mRNA splicing in the dorsolateral prefrontal cortex (DLFPC) of 450 subjects from two prospective cohort studies of aging. Hundreds of aberrant pre-mRNA splicing events are reproducibly associated with Alzheimer’s Disease (AD). We also generate a catalog of splicing quantitative trait loci (sQTL) effects in the human cortex: splicing of 3,198 genes is influenced by genetic variation. sQTLs are enriched among those variants influencing DNA methylation and histone acetylation. In assessing known AD loci, we report that altered splicing is the mechanism for the effects of the *PICALM, CLU,* and *PTK2B* susceptibility alleles. Further, we leverage our sQTL catalog to identify genes whose aberrant splicing is associated with AD and mediated by genetics. This transcriptome-wide association study identified 21 genes with significant associations, many of which are found in AD GWAS loci, but 8 are in novel AD loci, including *FUS,* which is a known amyotrophic lateral sclerosis (ALS) gene. This highlights an intriguing shared genetic architecture that is further elaborated by the convergence of old and new AD genes in autophagy-lysosomal-related pathways already implicated in AD and other neurodegenerative diseases. Overall, this study of the aging brain’s transcriptome provides evidence that dysregulation of mRNA splicing is a feature of AD and is, in some genetically-driven cases, causal.

## INTRODUCTION

Alternative splicing (AS) is an important post-transcriptional regulatory mechanism through which pre-mRNA molecules can produce multiple distinct mRNAs. AS affects over 95% of human genes^1^, contributing significantly to the functional diversity and complexity of proteins expressed in tissues^2^. AS is abundant in human nervous system tissues^3^ and contributes to phenotypic differences within and between individuals: at least 20% of disease-causing mutations may affect pre-mRNA splicing^4^. Mutations in RNA-binding proteins (RBPs) involved in AS regulation and aberrant AS have been linked to Amyotrophic lateral sclerosis (ALS)^5^ and Autism^6^. Further, disruptions in RNA metabolism, including mRNA splicing, are associated with age-related disorders, such as Frontotemporal lobar dementia (FTD)^7^, Parkinson’s disease^8^ and Alzheimer’s disease^9^,^10^. These studies have largely focused on alternative splicing of selected candidate genes, including the amyloid precursor protein (*APP*) ^8^ and microtubule associated protein *Tau* (*MAPT*)^8^,^9^,^11^. However, proteomics profiles of AD brains^12^ identified an increased aggregation of insoluble U1 snRNP, a small nuclear RNA (snRNA) component of the spliceosomal complex, suggesting that the core splicing machinery may be altered in AD. Apart from these studies, there have been few investigations of the possibility of more widespread splicing disruption affecting brain transcriptomes in AD^13^. However, a comprehensive study of *cis*-and *trans-* acting genetic factors that regulate alternative splicing in aging brains is lacking.

Over twenty-four genetic loci have now been associated with AD susceptibility by Genome-wide Association Studies (GWAS)^14^, and these AD variants are enriched for associations with gene expression levels in peripheral myeloid cells and often lie within *cis*-regulatory elements^15^. For example, we reported that one of these variants influences splicing of *CD33*^16^. Given the high abundance of alternative splicing in the brain, we hypothesized that other AD-associated genetic variants might also affect pre-mRNA splicing, possibly by disrupting efficient binding of splicing factors.

Here, by applying state-of-the-art analytic methods, we generated a comprehensive genome-wide map of splicing variation in the aging prefrontal cortex. We use this map to identify: (1) aberrant mRNA splicing events related to AD; (2) thousands of genetic variants influencing local mRNA splicing; (3) *trans* acting splicing factors that are involved in intron excision in brain; and (4) association of GWAS findings to specific genes that are likely to be causal in the etiology of AD. Overall, we deepen our understanding of genetic regulation in the aging brain’s transcriptome and provide a foundation for the formulation of mechanistic hypotheses in AD and other neurodegenerative diseases.

## RESULTS

### Aberrant mRNA splicing in AD and related-pathology

We deeply sequenced RNA from frozen dorsolateral prefrontal cortex (DLFPC) samples obtained at autopsy from 450 participants in either the Religious Order Study (ROS) or the Memory and Aging Project (MAP), two prospective cohort studies of aging that include brain donation. All subjects were without known dementia at study entry. During the study, some subjects experienced cognitive decline, and, at autopsy, they displayed a range of amyloid-β and *Tau* pathology, with 60% of subjects having a pathologic diagnosis of AD^17^,^18^ (**Supplementary Table 1**).

Following alignment and quantification of RNA-Seq reads, LeafCutter^19^,^20^ was applied to estimate “percent spliced in” values (PSI, Ψ) for local alternative splicing events (**Fig. 1**). LeafCutter detects splicing variation using those reads that span splice junctions. We identified 53,251 alternatively spliced intronic excision clusters in 16,557 genes. To identify aberrant splicing events, we analyzed the association between the PSI of each intron excision event and a pathologic diagnosis of AD or quantitative measures of AD neuropathology including neuritic plaques (NP), neurofibrillary tangles (NFT), and amyloid-β burden, while accounting for confounding factors (**Online Methods**). At False Discovery Rate (FDR) < 0.10 we identified a total of 303 differentially spliced introns in 224 genes associated with different AD neuropathologies including 13 with NP, 82 with amyloid, and 234 with NFT (**Supplementary Table 2).** A heat map of the top differentially spliced introns associated with NFT is shown in **Fig. 2a**. On average, these differentially excised introns explain ∽2-13% of total variation in different neuropathologies after accounting for biological (age and sex) and technical covariates (**Fig. 2b; Supplementary Fig. 1**).

**Figure 1:**
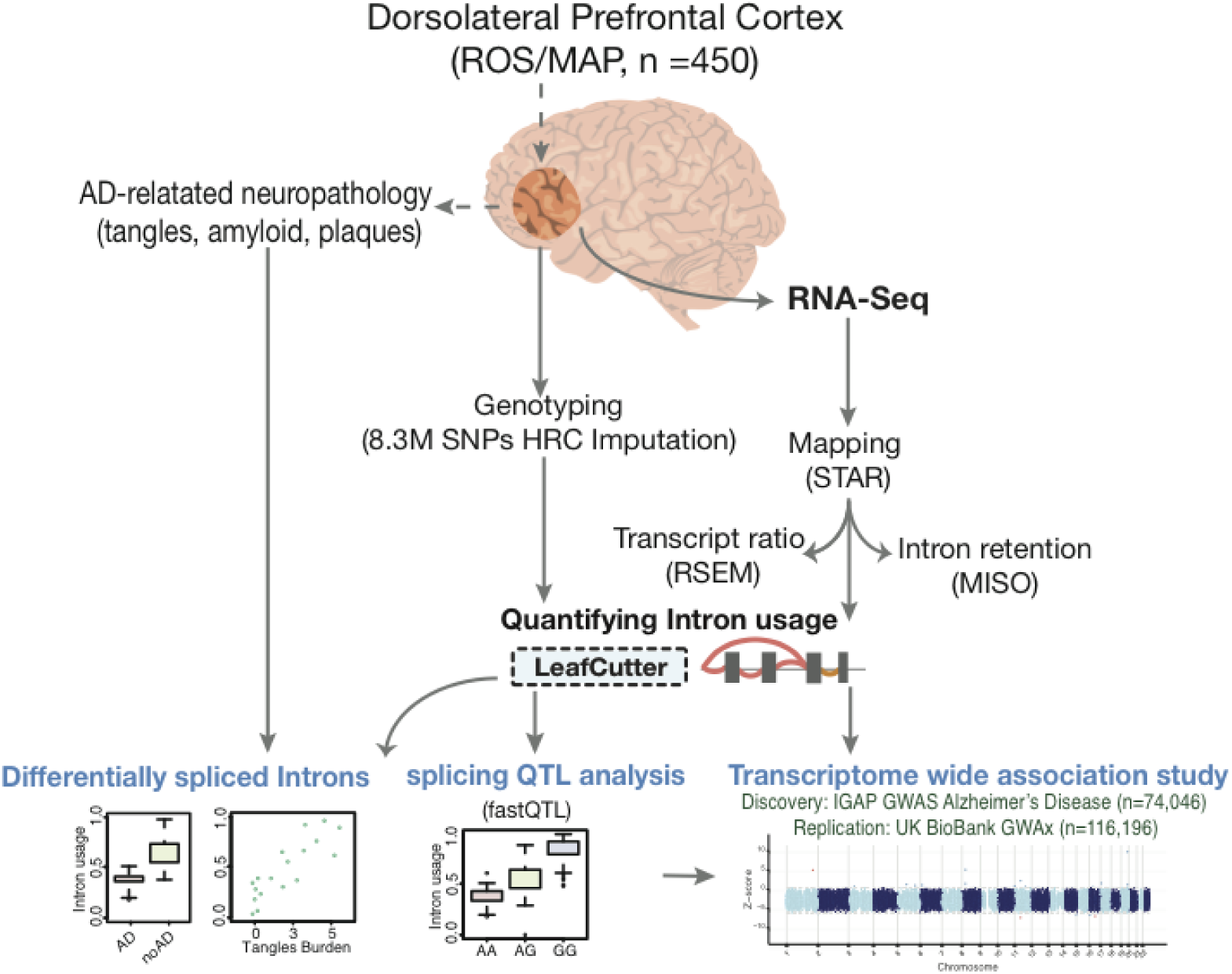
Overview of the study. RNA was sequenced from the gray matter of the dorsal lateral prefrontal cortex (DLPFC) of 542 samples (461 remained after QC and matching for genotype data) from the ROS/MAP cohort. RNA-Seq data were processed, aligned and quantified by our parallelized pipeline. The intronic usage ratios for each cluster were then computed using LeafCutter^19^,^20^, standardized (across individuals) and quantile normalized. The intronic usage ratios were used for differential splicing analysis, for calling splicing QTLs, and for transcriptome-wide association studies (TWAS). TWAS was performed on summary statistics from IGAP AD GWAS of 74,046 individuals^37^.

**Figure 2:**
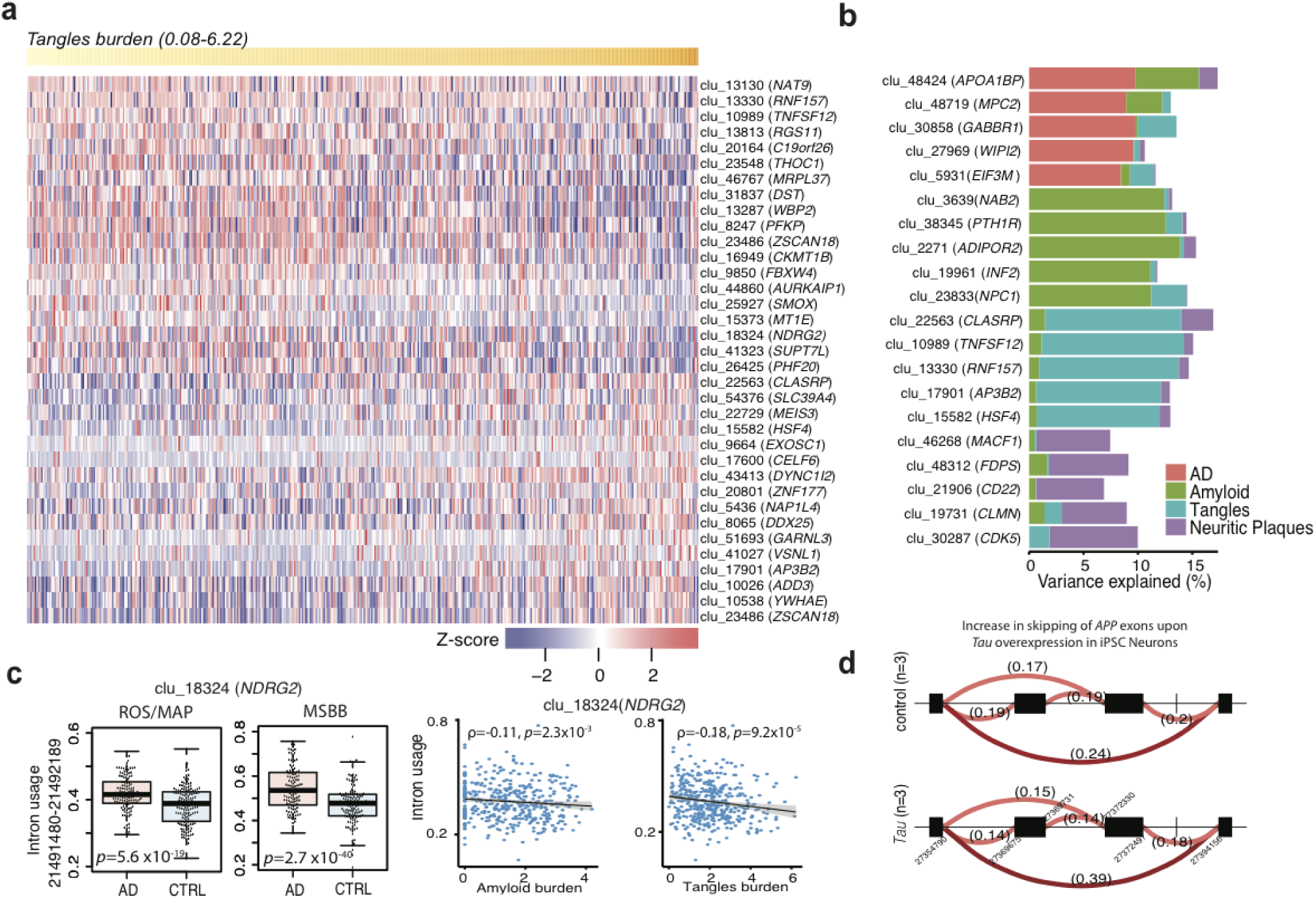
Differential splicing analysis in relation to AD diagnosis and AD neuropathology. (a) Heat map of top 35 differently excised intron association with burden of tangles in ROSMAP. Each column is one subject, who are ordered by their tangles burden (yellow row at the top of the panel). The association’s Z-score strength and direction are denoted using the key at the bottom of the panel. (b) Variance explained (%) of top 5 differently excised introns association for four different traits. (c) The left two panels present the mean and distribution of intron usage for differently excised introns in *NDRG2* in relation to a clinical diagnosis of AD in ROSMAP and in MSBB. The right two panels display the association of amyloid or tangle burden to intron usage in *NDRG2*. (d) Differentially excised intron in *APP* upon *Tau* overexpression in iPSC Neurons.

To test for association with the clinical diagnosis of AD, we used Leafcutter^19^,^20^ to identify differentially spliced introns by jointly modeling intron clusters using a Dirichlet-multinomial GLM (**Online Methods**). At a Bonferroni-corrected *P* < 0.05, we identified a total of 87 intron clusters (corresponding to 84 genes) that displayed altered splicing in relation to AD (**Supplementary Table 3**). For example, the most significant differentially excised intron (chr10: 3147351-3147585) in the gene *PFKP*, phosphofructokinase: the frequency of this event was associated with AD (P < 4.9× 10^−24^) and all pathologic measures tested in this study. Similarly, the next most differentially excised intron (chr14: 21490656-21491400) associated with AD is found in the alpha/beta-hydrolase fold protein gene *NDRG* family member 2 (*NDRG2*) (*P* < 5.6× 10^−19^) and is also associated with measures of both amyloid and *Tau* pathology (**Fig. 2c**). Differential splicing of both *PFKP* and *NDRG2* in human brains has been previously shown to be associated with AD pathogenesis^21^,^22^, offering a measure of replication. Other genes with differentially excised introns associated with AD include *APP* (*P* < 1.6× 10^−3^) and genes in known AD GWAS loci including *PICALM* (*P* < 0.02) and *CLU* (*P* < 3.2 × 10^−4^). Next, to assess the robustness of our results, we performed a replication analysis using RNA-Seq data from the Mount Sinai Brain Bank (MSSB)^23^ involving 301 samples from AD and control brains (see **Supplementary Note**). Of the 84 genes with differentially spliced intron clusters in ROSMAP, 52 (including *APP*, *PFKP, and NDRG2*) were significant at a Bonferroni-corrected *P* < 0.05 thresholds in the MSBB data (**Fig. 2c; Supplementary Table 4**). This constitutes an independent replication of specific, aberrant splicing alterations in AD brains. Finally, to further validate and explore the mechanism of our observations, we analyzed RNA-Seq data derived from control iPSC-derived neurons (iN) and iN overexpressing *Tau*: differential intron excision was noted at *APP* (*P* < 4.9 × 10^−6^) and *NDRG2* (*P* < 0.006) in this model system (**Fig. 2d**). These data suggest that *tau* accumulation in neurons – at a stage in which neurons are accumulating phospho-*tau* but are not apoptotic-is sufficient to induce splicing alterations; this *in vitro* validation of disease-related splicing changes suggests that (1) altered splicing is not related to confounding factors relating to autopsy or the agonal state and (2) has specific target RNAs that can be modeled *in vitro*.

### Genetic effects on pre-mRNA splicing in aging brains

We next performed a splicing QTL (sQTL) study to identify local genetic effects that drive variation in RNA splicing in the DLFPC. First, we assessed the splice events from the LeafCutter algorithm (**Fig. 1**); 30% of these 54,463 intron excision clusters are novel splicing events, not previously reported in other sQTL studies. The PSI values were adjusted for known and hidden factors (15 principal components) and then fit to imputed SNP data using an additive linear model implemented in fastQTL^24^ (**Online Methods; Supplementary Fig. 2**). At FDR < 0.05, we found 9,028 sQTLs in 3,006 genes (**Supplementary Table 5**). Over 60% of these sQTLs involve changes in cassette splicing (simple or complex exon skipping events), followed by 5’/3’ exon extension (23%), and alternative upstream or downstream exon usage (17%). As expected, splicing was most strongly affected by variants in the splice region itself (59.8%): 20.2% of variants are mapped to splice acceptor sites and 16.4% to splice donor sites. The remaining (23.2%) mapped to other splice regions or are found within an intron **(Supplementary Fig. 3).** Further, sQTLs are mapped to distinct regulatory features as defined by 15 chromatin states in DLPFC^25^: sQTLs were significantly enriched in actively transcribed regions and enhancers. They are depleted in repressed chromatin marked with polycomb, heterochromatin, and quiescent regions (**Fig. 3a**).

**Figure 3:**
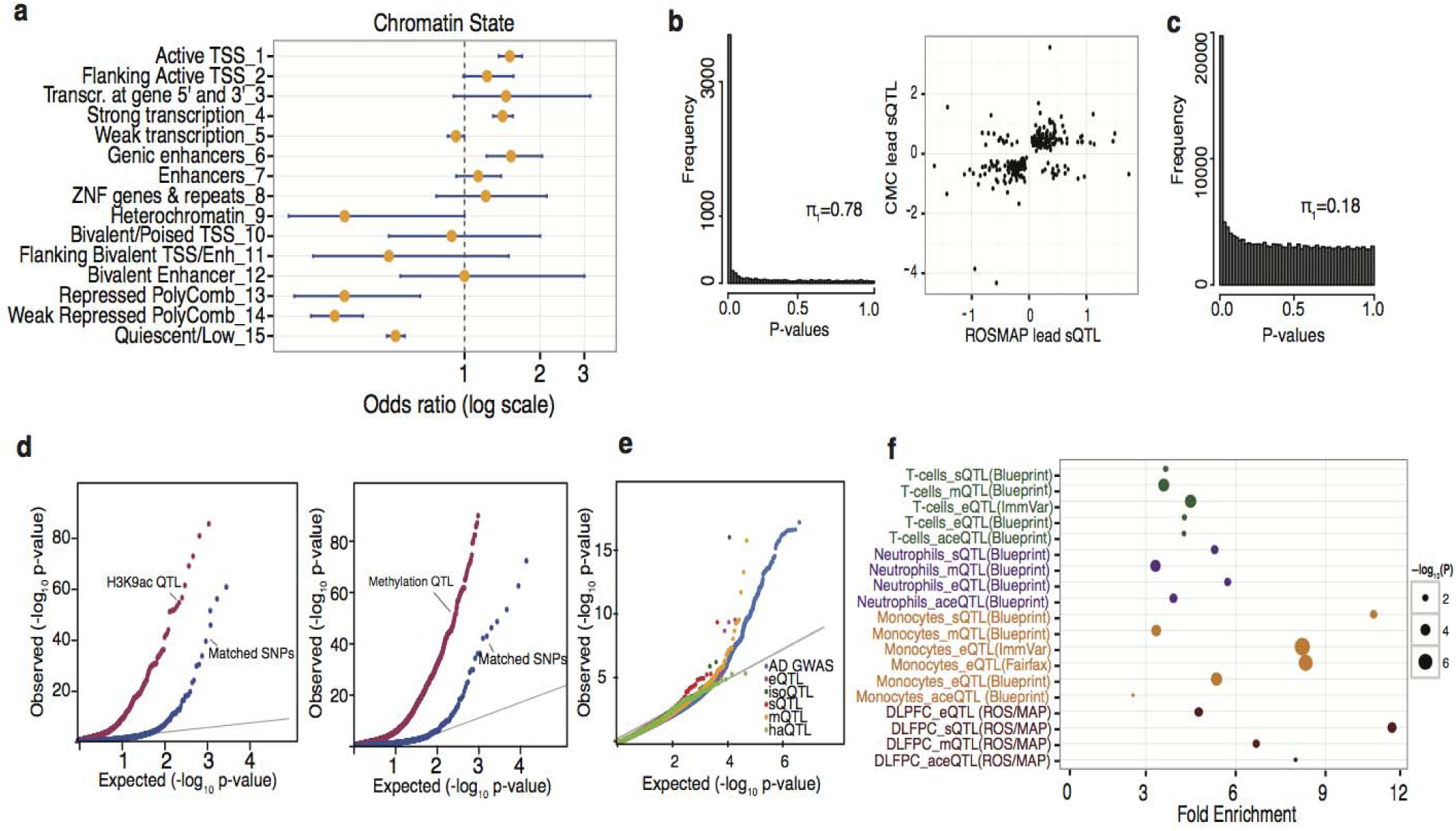
Enrichment of splicing QTLs in epigenomic marks and in AD GWAS. (a) Splicing QTLs are enriched in regions (or chromatin states) associated with active transcription and genic enhancers, and they are depleted in polycomb regions that are transcriptionally repressed in the DLPFC. (b) Left: P-value distribution of ROSMAP sQTLs that are significant in CMC (FDR < 0.05). The majority (78%) of sQTLs in ROSMAP are also discovered in CMC. Right: The direction of effect is consistent for the majority (93%) of the significant (FDR 0.05) lead sQTLs in CMC and in ROSMAP. (c) P-value distribution of ROSMAP eQTLs that are significant sQTLs (FDR < 0.05). (d) SNPs that drive QTLs in H3K9ac and DNA methylation data in the same ROSMAP brains are more likely to be sQTLs than matched SNPs within H3K9ac domains (left) and near DNA methylated CG (right). (e) QQ-plot for AD GWAS suggests that sQTLs are enriched among AD GWAS (IGAP study^37^) compared to other types of QTLs. (f) Fold-enrichment of AD GWAS SNPs (*P* < 10^-5^) among QTL SNPs driving variation in gene expression, splicing, histone acetylation, and DNA methylation in primary monocytes^15^,^29^,^64^, T-cells^15^,^29^, or DLFPC^27^.

To assess the extent of sQTL replication, we compared our sQTLs to the recently published dataset from the CommonMind Consortium (CMC), consisting of DLFPC profiles from 258 persons with schizophrenia and 279 control subjects^26^ (see Supplementary Notes). Our sQTLs yield a Storey’s Π_1_ = 0.78 in the CMC data, suggesting substantial sharing of sQTLs between these two different brain collections (**Fig. 3b**). Moreover, 93% of sQTLs showed the same direction of effect (**Fig. 3b**). The fraction of sQTLs that are novel deserve further evaluation to assess the extent to which they may be context-specific given that the average age at death of our participants is 88 years.

In agreement with recent findings in lymphoblastoid cell lines (LCLs)^20^, we found that a majority of sQTLs act independent of gene expression effect, as evident by the low degree of sharing between sQTLs and eQTLs27 from the same brains (Π_1_ = 0.18) (**Fig. 3c**). Of the 9,045 lead sQTL SNPs, only 42 are also a lead eQTL, suggesting that a substantial fraction of sQTLs are unique and are not detected by standard eQTL analysis.

To further understand the mechanisms underlying sQTLs, we assessed the overlap of sQTLs with SNPs influencing epigenomic marks (xQTLs) such as DNA methylation (mQTL) and histone H3 acetylation on lysine 9 (H3K9Ac, haQTL)^27^ that are available from the same DLPFC samples (**Online Methods**). Indeed, we found that such xQTLs27 are significantly enriched among sQTLs when compared to randomly selected, matched SNPs (Kolmogorov–Smirnov test *P* < 0.001 (**Fig. 3d)**: of the lead sQTL, 9% (578) and 19% (1246) were also associated with haQTL and mQTL, respectively, suggesting extensive genetic co-influences on splicing, methylation, and histone modifications. Finally, we found significant sharing of sQTL SNP among SNP that also influence histone (*Π*_1_ = 0.74) or methylation (*Π*_1_ = 0.82). These overlaps suggest a contribution of epigenomic regulation in directing splicing.

Given prior reports^20^,^28^, we evaluated whether our sQTLs from the aging brain were enriched for AD susceptibility variants (**Figs. 3e and 3f**). We also assessed enrichment of AD SNPs (*P* < 1× 10^−5^) in splicing, methylation or expression QTLs from DLFPC^27^, monocytes^15^,^29^, neutrophils^29^ and T-cells^15^,^29^. Using an enrichment method that uses permutation testing (matching for MAF, distance to TSS, and a number of LD proxies), we found that DLFPC sQTLs are more likely to be enriched among AD GWAS SNPs (*P* <10^−5^), followed by sQTL and eQTL from monocytes (**Fig. 3f**). These findings suggest the important role of RNA splicing on variation in AD susceptibility, the prominent role of myeloid cells in AD susceptibility^15^ but also the fact that a number of AD variants have mechanisms that may be mediated through non-myeloid effects.

Some of these effects of AD variants on splicing are known, such as the 8-fold increase in full-length *CD33* isoform^16^,^30^ and the *SPI1* functional consequence^31^, but several of these-in *CLU*, *PICALM,* and *PTK2B*-have not been previously reported (**Supplementary Table 6**). For example, the *PTK2B* risk allele leads to increased skipping of exon 31 (chr8: 27308560-27308595) which contains a coding part of the gene (**Fig. 5d**). These results delineate the initial events along the cascade of functional consequences for these three AD variants and provide important mechanistic insights into their development as potential therapeutic targets.

### Splicing regulators associated with alternative splicing in DLFPC

Splicing of pre-mRNA is catalyzed by a large ribonucleoprotein complex called the spliceosome, which consists of five small nuclear RNAs (snRNA) and numerous splicing factors^32^. To identify brain splicing factors that regulate sQTL events in *trans*, we evaluated whether the lead sQTL SNPs identified in our study are enriched in RNA binding protein (RBP) binding sites using publicly available cross-linking immunoprecipitation (CLIP)-Seq datasets from 76 RBPs in CLIPdb^33^. We found that binding targets of 18 RBPs are significantly enriched among lead sQTLs (**Fig. 4a**). The most enriched RBP is *PTBP1* (Fisher’s exact P < 0.006), followed by *HNRNPC*, *CPSF7*, and *ELAVL1*. Notably, the enrichment for neuronal *ELAVL1* RBP target sites is consistent with a recent report that, upon neuronal *ELAVL1* depletion, *BIN1* and *PICALM* transcripts were found to have lower exon inclusion for those sites in which *ELAVL* binding sites directly overlapped with SNPs associated with AD^34^.

**Figure 4:**
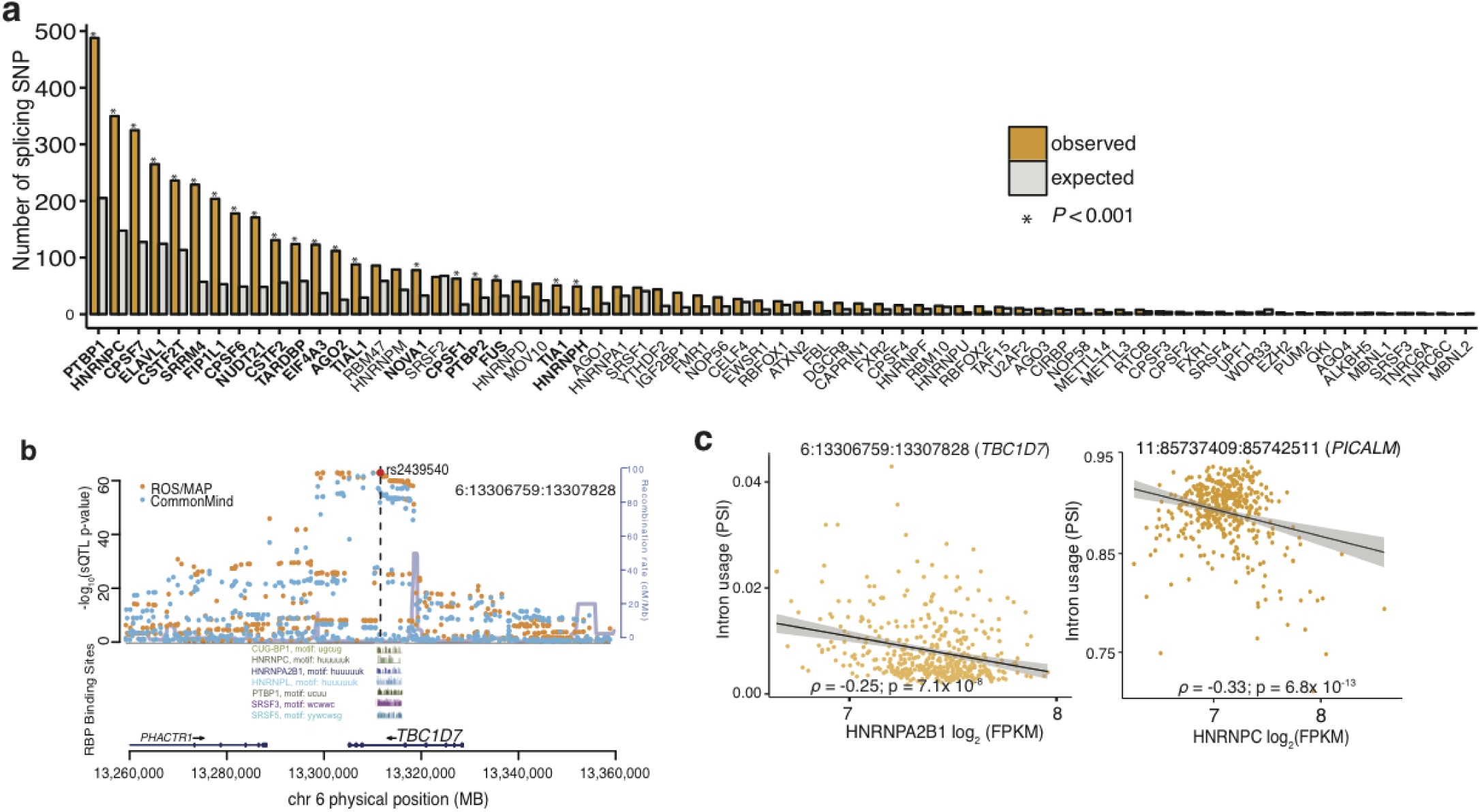
Enrichment of RNA-binding protein (RBP) binding sites among sQTLs. (a) RBP enrichment (expected vs. observed) among the lead sQTLs. Significant RBSs are in bold and shown with an “*”. (b) Regional plot of sQTL results for SNPs in the vicinity of *TBC1D7* (6:13306759:13307828). SNPs driving splicing QTLs for *TBC1D7* overlap CLIP binding sites (from CLIPdb^33^) for several splicing factors. The top SNP (rs2439540, red color) overlaps motifs for a number of RBPs. Splicing QTL results are highly consistent between ROSMAP (orange) and CommonMind (blue) data. (c) Association of *hnRNPA2B1* (left) and *hnRNPC* (right) gene expression levels with differential intron usage in *TBC1D7* (left) and in *PICALM* (right).

On the other hand, we also observed significant enrichment for the lead sQTL SNPs within the binding sites for a number of heterogeneous nuclear ribonucleoproteins (hnRNP) including hnRNP C (*P* < 0.009). Further, we find that the expression levels of hnRNP splicing factors are correlated with intronic excision levels of hundreds of genes, many of which are in AD susceptibility loci including *BIN1, PICALM, APP, and CLU* **(Supplementary Figs. 4 and 5).** The hnRNP C factor has been linked to AD in previous studies, including in a recent biochemical study reporting the translational regulation of *APP* mRNA by hnRNP C^35^. This observation goes towards the mechanism of the sQTL: consistent with the assumption that, altering the sequence of a binding site changes the likelihood that a splicing event occurs *in vivo*. In one example of a sQTL affecting intron usage, a *TBC1D7* (one of the new AD genes described in a later section**; Figs. 6a and 6b**) SNP is found within CLIP-defined binding sites for hnRNP C as well as other RBPs (**Fig. 4b**). Thus, incorporating RBP binding sites as a functional annotation allows for improving our accuracy in selecting plausible causal variants that may disrupt binding of splicing factors to cause the alternative-splicing event. Further biochemical studies will be required to understand the full regulatory program that orchestrates the disease-related splicing changes.

### Transcriptome-wide association studies prioritizes AD genes in endocytosis and autophagy-lysosomal pathways

To identify genes whose mRNA expression or alternative splicing is associated with AD and mediated by genetic variation, we performed two Transcriptome-wide association studies (TWAS)^36^ by using either the ROSMAP expression data or its intronic excision levels as reference panels to re-analyze summary level data from the International Genomics of Alzheimer’s Project (IGAP) AD GWAS that includes data from 79,845 individuals^37^. Using the reference data, this method infers expression levels into the IGAP summary statistics, performs a case/control analysis of the imputed expression or splicing data, and generates a joint association statistic that integrates the extent of genetic and expression or splicing association for a given gene. A total of 4,746 genes and 15,013 differentially spliced introns could be analyzed, and we identified 21 genes at FDR < 0.05 whose imputed gene expression or intronic excision levels were significantly associated with AD status (**Fig. 5a; Supplementary Table 6**). Among these, there were genes in known AD loci including *SPI1, CR1, PTK2B*, *CLU*, *MTCH2,* and *PICALM*. These results help to pinpoint the likely gene that is the target of the known susceptibility variant in each locus, particularly at the *MTCH2* locus in which the functional consequence of the risk allele was unclear. However, the new AD genes are even more interesting, and 8 of these associations are found in loci that harbored only suggestive evidence of association in the IGAP study. These genes include *AP2A1*, *AP2A2*, *FUS*, *MAP1B, TBC1D7* and others that are now significant at a threshold adjusted for genome-wide testing and therefore help to prioritize the long list of suggestive IGAP associations (**Figs. 5a, 6a, and 6b**; **Supplementary Figs. 6-14**). *FUS* is particularly intriguing since mutations in *FUS* have been previously linked to ALS^38^; this suggests that there may be some shared genetic susceptibility between these two neurodegenerative diseases that have not been appreciated previously.

**Figure 5:**
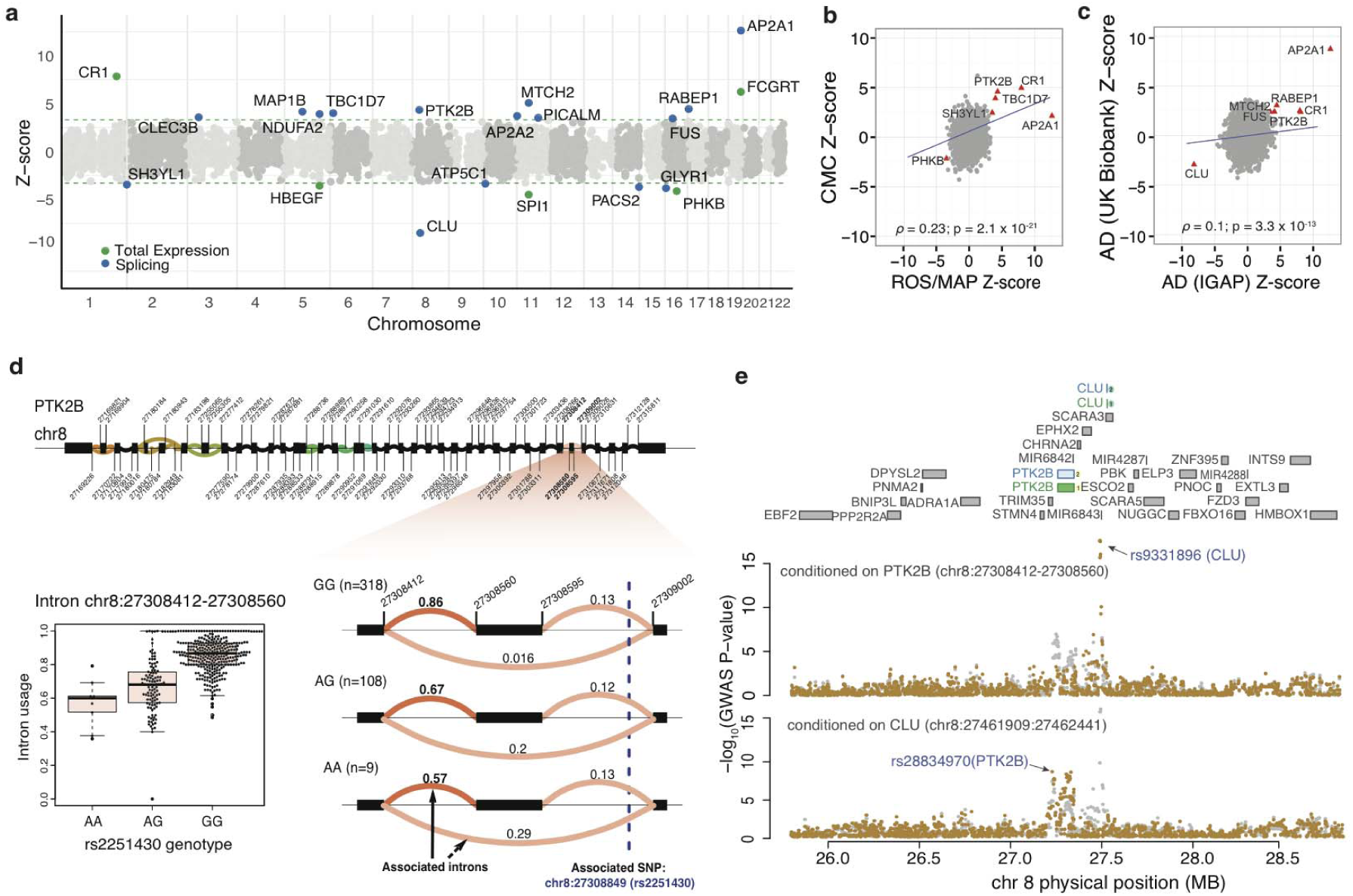
Transcriptome-wide association study of Alzheimer’s Disease. (a) Transcriptome-wide results using the IGAP AD GWAS summary statistics based on data from 74,046 individuals; each dot is one gene. The dotted green line denotes the threshold of significance (FDR 0.05). Genes for which there is evidence of significant differential intron usage are highlighted in blue. In green, we highlight those genes where the TWAS using total gene expression results are significant. (b) Replication of ROSMAP TWAS in CMC DLFPC data. The red triangles denote genes where the replication analysis is significant. (c) Replication of IGAP AD TWAS using the UK BioBank AD GWAS based on an independent set of subjects. (d) *PTK2B* gene structure (top): clusters of differential splicing events are noted with the colored curves. The panel then zooms to highlight differential intronic usage for chr8:27308412-27308560 stratified by rs2251430 genotypes (right). On the left, we show the same data use a box plot. (e) Conditional analysis of IGAP AD GWAS results for two splicing effects for *PTK2B* and *CLU* in AD GWAS data. As noted in the top aspect of the panel, these two AD genes are located close to one another. The intronic excision events for *PTK2B* and *CLU* are present in both ROSMAP (blue) and in CMC (green) dataset. When the AD GWAS is conditioned on the *PTK2B* (chr8:27308412-27308560) splicing effect, the *CLU* effect remained significant, demonstrating its independence from the PTK2B association (superior locus plot where each dot is one SNP in the genomic segment under evaluation). The reciprocal analysis conditioning on the *CLU* (chr8:27461909:2746244) effect, the *PTK2B* association remained significant.

To replicate these results, we first assessed whether using the expression imputation model built using the CMC dataset^26^ that was deployed in IGAP AD GWAS yields significant results. We focused on the 21 significantly associated genes above: five genes (*CR1*, *PTK2B, CLU, TBC1D7, and AP2A2*) replicated at FDR < 0.05 and two genes (*MTCH2* and *PICALM*) were nominally significant at *P* < 0.05 with the expression and splicing inference from CMC (**Fig. 5b**). The directions of effect for all six associations were consistent in both datasets (**Fig. 5b**). Thus, we see robust replication, and our results are not due to the unique properties of the ROSMAP dataset. Second, we used the UK BioBank (UKBB) AD GWAS by proxy^39^ to replicate the IGAP TWAS results. We note that, despite analyzing data from 116,196 subjects, the UKBB AD GWAS is underpowered since the GWAS does not use AD cases but, rather, subjects who have a first-degree relative with AD as “cases”. Nevertheless, we were able to replicate (at nominal *P* < 0.05) seven of our IGAP TWAS associations in the UKBB TWAS (**Fig. 5c**). These two complementary replication efforts demonstrate the robustness of our results. Finally, we performed a TWAS using the summary statistics of a meta-analysis of IGAP and UKBB GWAS, and identified three additional genes (*ABCA7*, *RHBDF1*, and *VPS53*) that meet a genome-wide significant threshold in the meta-analysis (**Supplementary Table 7**), with ABCA7 being one of the well-validated AD loci.

Most of the TWAS associations are the result of differential intron usage, suggesting the importance of pre-mRNA splicing in AD (**Fig. 5a**). An example of TWAS association with intron usage at *PTK2B*, a known AD susceptibility locus, is shown in **Fig. 5d**. We often observed multiple TWAS-associated genes in the same locus, likely due to co-expression of genes in close physical proximity or allelic heterogeneity within the susceptibility locus^40^. To account for multiple associations in the same locus, we applied conditional and joint association methods that rely on summary statistics^40^,^41^ to identify genes that had significant TWAS associations when analyzed jointly (**Online Methods; Figs. 5e and 6b**; **Supplementary Figs. 6-14**). A region with multiple TWAS association includes the *PTK2B/CLU* locus, which shows independent co-localized association for both GWAS^37^ and splicing effects (**Fig. 5e**).

Refining known associations is important to translate results into functional studies, but the newly validated AD genes offer new insights into AD: we used the GeNets (http://apps.broadinstitute.org/genets) to evaluate the connectivity of our new AD genes with the network of known AD susceptibility genes that are interconnected by protein-protein interaction (PPI)^42^. These new and known AD susceptibility genes are directly connected (i.e., they form shared ‘communities’) (*P* < 0.006) (**Fig. 6c**). Further, this joint network is enriched for endocytosis pathways (*P* < 0.0002), highlighting the existing narrative of endocytosis pathways being preferentially targeted in AD. More interesting is the enrichment for the autophagy-lysosomal related pathway (*P* < 0.003) (**Fig. 6c**). The genes in the autophagy-lysosomal related pathway (*AP2A2, AP2A1,* and *MAP1B*) form a statistically significant *P* < 4.3× 10^−4^) PPI sub-network with known AD genes (*PTK2B*, *PICALM* and *BIN1*) (**Fig. 6d**). Protein degradation pathways have been implicated previously in ALS^43^ and to a limited extent in AD^44^. Overall, these PPI analyses suggest that our new TWAS-derived genes are not a random set of genes but are part of an AD network.

**Figure 6:**
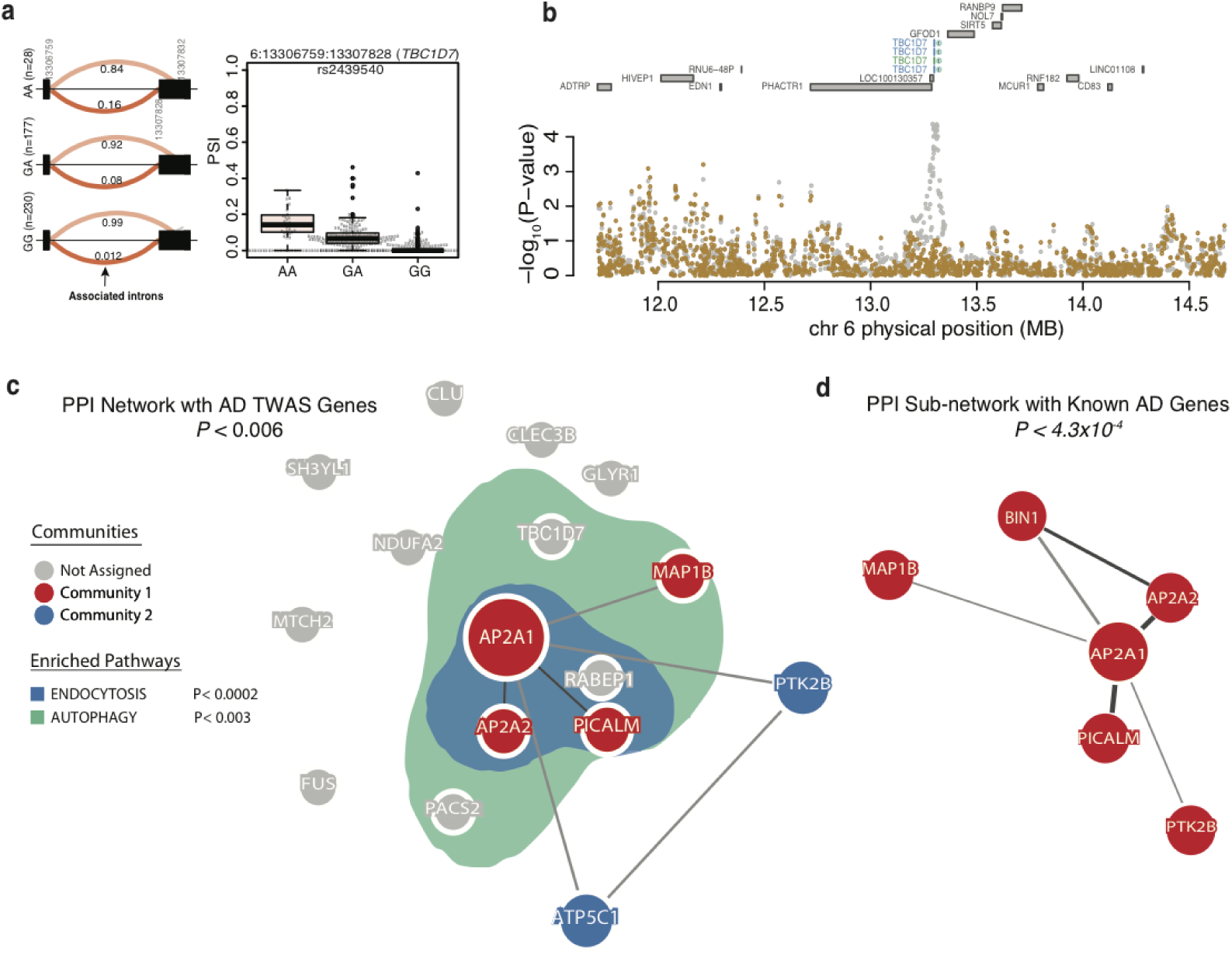
TWAS prioritizes AD genes in endocytosis and autophagy-related pathway. (a) Differential intronic usage for 6: 13306759:13307828 (*TBC1D7*) stratified by rs2439540 genotypes (left). Box plot for the same data (right). (b) Regional plot showing the AD IGAP P-values in *TBC1D7* locus. Two intronic excision events at *TBC1D7* are present in both ROSMAP (blue) and in CMC (green) dataset. The AD GWAS effect is mostly explained by intronic usage of 6:13306759:13307828. The AD GWAS at *TBC1D7* is suggestive in the original IGAP study (p<10^-5^). (c) The product of three of the novel AD genes (*AP2A2, AP2A1*, and *MAP1B*) are members of the same PPI network (*P* < 0.006). The genes in this network and others not in the network (i.e., *TBC1D7, PACS2*, and *RABEP1*) are significantly enriched in genes annotated as being involved in endocytosis (blue; *P* < 0.0002) and autophagy-related pathways (green; *P* < 0.003). (d) The novel AD genes (*AP2A2, AP2A1,* and *MAP1B*) form a significant PPI sub-network (*P* < 4.3 × 10^−4^) with known AD genes (i.e., *PICALM, BIN1*, and *PTK2B*).

## DISCUSSION

In this study, we directly examined alternative splicing events in a large dataset of aging brains, which led to both the observation that specific alternative splicing events are reproducibly associated with AD and the functional dissection of genetic associations to AD. Our replication efforts demonstrate that the observed AD-related perturbations in splicing are not simply due to spliceosomal failure. Further, our *in vitro* model of *tau* overexpression in iPSC-derived neurons shows that perturbation of *MAPT* is sufficient to cause these disease-related splicing changes that are observed in the human cortex at autopsy. Finally, since these neurons are functionally normally, we now know that these splicing changes occur very early in the series of molecular events that are caused by perturbation in *MAPT* expression.

We combined this splicing map of the aging brain with common genetic variants and cataloged the genetic architecture controlling local splicing events. These analyses revealed a preponderance of brain and myeloid splicing events as being the functional consequence of AD susceptibility alleles, which connects with the broader AD-related splicing changes to highlight the role of altered RNA maturation as playing a key role in this neurodegenerative disease (**Figs. 3f and 5a; Supplementary Table 6**).

To address the issue of causality in results derived from our cross-sectional brain data, we used the powerful TWAS approach, which leverages our splicing map and common genetic variants to test the hypothesis that the effect of such variants in AD is mediated, by altering splicing levels. These analyses confirmed many of the known AD genes (i.e., *CLU* and *PTK2B*), which supports the role of regulation of splicing levels as key mechanisms in certain AD loci, but also found several new AD loci: *TBC1D7, AP2A1, AP2A2*, *FUS,* and *MAP1B* (**Figs. 5a and 6b; Supplementary Figs. 6,8, and 12)**. These new genes reinforce the association of the Clathrin/AP2 adaptor complex with AD susceptibility^45^. Both *AP2A2* and *AP2A1*, which are components of the AP2 adaptor complex that serves as a cargo receptor, selectively sorting membrane proteins involved in receptor-mediated endocytosis^46^. The AP2 complex and PICALM interact with APP, directing it to degradation and autophagy^46^.

Our study also offers insights for several well-known AD loci in which the gene was known but the functional mechanism remained unclear. Similar to our work in *CD33*^16^, the careful analysis of these cortical data highlights a specific splicing mechanism for the AD risk alleles at *CLU, PICALM,* and *PTK2B*. All three are complex proteins with a large number of exons, so our results prioritize specific domains in these proteins as harboring the functional domain that influences AD risk. These domains will be critical in beginning to assemble a protein:protein interaction scaffold for AD susceptibility that goes beyond repurposing existing databases of interactions that are assessed in GeNets. Further, our analyses of RBP involved in splicing regulation of AD susceptibility genes including *PICALM* and RNA binding site analysis of *HNRNPC* (**Fig. 4c; Supplementary Fig. 5**) and *ELAVL* helps to prioritize the putative causal variant and to elaborate the series of events upstream of the susceptibility variant that enable its expression. Thus, our catalog of splicing variants made available with this study provides a starting point for further focused molecular and biochemical experimental validation to fully elucidate the role of these splicing variants in the etiology of AD.

This transcriptome-wide reference map of RNA splicing in the aging cortex is a new resource that highlights strong effects of neuropathology and genetic variation on splicing. It will be useful in annotating the results of genetic and epigenomic studies of neurologic and psychiatric diseases, but it has an immediate impact in (1) identifying the functional consequences of several AD susceptibility alleles, (2) extending the list of loci involved in AD, and (3) implicating the protein degradation machinery in the pathology of AD.

## METHODS

### Study Cohorts

#### Religious Orders Study (ROS)

From January 1994 through June of 2010, 1,148 persons agreed to annual detailed clinical evaluation and brain donation at the time of death. Of these, 1,139 have completed their baseline clinical evaluation: 68.9% were women; 88.0% were white, non-Hispanic; their mean age was 75.6 years; and mean education was 18.1 years. There were 287 cases of incident dementia and 273 cases of incident AD with or without a coexisting condition. Details of the clinical and pathologic methods have been previously reported ^17^.

#### Memory and Aging Project (MAP)

From October 1997 through June 2010, 1,403 persons agreed to annual detailed clinical evaluation and donation of the brain, spinal cord, nerve, and muscle at the time of death. Of these, 1,372 completed their baseline clinical evaluation: 72.7% were women; 86.9% were white, non-Hispanic; their mean age was 80.0 years; and mean education was 14.3 years with 34.0% with 12 or fewer years of education. There were 250 cases of incident dementia and 238 cases of incident AD with or without a coexisting condition. Details of the clinical and pathologic methods have been previously reported ^47^. To avoid population stratification artifacts in the genetic analyses, the study was limited to non-Latino whites.

See Supplementary Notes for the details of CommonMind Consortium (CMC) and Mount Sinai Brain Bank (MSBB) datasets.

### Data acquisition, quality control, and normalization

#### Genotyping

DNA from ROS and MAP subjects was extracted from whole blood, lymphocytes or frozen post-mortem brain tissue and genotyped on the Affymetrix GeneChip 6.0 platform at the Broad Institute’s Center for Genotyping. Only self-declared non-Latino Caucasians were genotyped to minimize population heterogeneity. PLINK software^48^ was used to implement our QC pipeline. We applied standard QC measures for subjects (genotype success rate >95%, genotype-derived gender concordant with reported gender, excess inter/intra-heterozygosity) and for single nucleotide polymorphisms (SNPs) (HWE P > 0.001; MAF > 0.01, genotype call rate > 0.95; misshap test > 1× 10^−9^) to these data. Subsequently, EIGENSTRAT^49^ was used to identify and remove population outliers using default parameters. Imputation was performed using Michigan Imputation Server with Minimac3^50^ using Haplotype Reference Consortium (HRC version r1.1, 2016)^51^ panel consisting of 64,940 haplotypes of predominantly European ancestry. Imputation filtering of r^2^ > 0.3 was used for quality control. After QC, 450 individuals and 8,383,662 genotyped or imputed markers were used for sQTL analysis.

#### RNA-Seq data

RNA was sequenced from the gray matter of dorsal lateral prefrontal cortex (DLPFC) of 542 samples, corresponding to 540 unique brains. These samples were extracted using Qiagen’s miRNeasey mini kit and the RNase free DNase Set. RNA was quantified using Nanodrop. The quality of RNA was evaluated by the Agilent Bioanalyzer. All samples were chosen to pass two initial quality filters: RNA integrity (RIN) score >5 and quantity threshold of 5 μg (and were selected from a larger set of 724 samples). RNA-Seq library preparation was performed using the strand specific dUTP method14 with poly-A selection. Sequencing was performed on the Illumina HiSeq with 101bp paired-end reads and achieved coverage of 150M reads of the first 12 samples. These 12 samples served as a deep coverage reference and included 2 males and 2 females of non-impaired, mild cognitive impaired, and Alzheimer’s cases. The remaining samples were sequenced with target coverage of 50M reads; the mean coverage for the samples passing QC is 95 million reads (median 90 million reads). The libraries were constructed and pooled according to the RIN scores such that similar RIN scores would be pooled together. Varying RIN scores result in a larger spread of insert sizes during library construction and leads to uneven coverage distribution throughout the pool.

The RNA-Seq data were processed by a parallelized pipeline. This pipeline includes trimming the beginning and ending bases from each read, identifying and trimming adapter sequences from reads, detecting and removing rRNA reads, and aligning reads to reference genome. Specifically, RNA-Seq reads in FASTQ format were inspected using FASTQC program. Barcode and adapter contamination, low-quality regions (8bp at beginning and 7bp at ending of each FASTQ reads) were trimmed using FASTX-toolkit. To remove rRNA contamination, we aligned trimmed reads to rRNA reference (rRNA genes were downloaded from UCSC genome browser selecting the RepeatMask table) by BWA then extracted only paired unmapped reads for transcriptome alignment. STAR (v2.5)^52^ (was used to align reads to the transcriptome reference, and RSEM (v1.3.0)^53^ was used to estimate expression levels for all transcripts. To quantify the contribution of experimental and other confounding factors to the overall expression profiles, we used the COMBAT algorithm^54^ to account for the effect of batch and linear regression to remove the effects of RIN, post-mortem interval (PMI), sequencing depth, study index (ROS sample or MAP sample), genotyping PCs, age at death, and sex. Finally, only highly expressed genes were kept (mean expression >2 log_2_-FPKM), resulting in 13,484 expressed genes for eQTL analysis. The details for *cis*-eQTL analysis are in Ng et al.^27^.

#### Intron usage mapping and quantification

We used LeafCutter^19^,^20^ to obtain clusters of variably spliced introns. Leafcutter allows the identification of splicing events without relying on existing annotations, which are typically incomplete, especially in the setting of large genes or individual/population-specific isoforms. Leafcutter defines “clusters” of introns that represent alternative splicing choices. To do this, it first groups together overlapping introns (defined by spliced reads). For each of these groups, Leafcutter constructs a graph where nodes are introns and edges represent overlapping introns. The connected components of this graph define the intron clusters. Singleton nodes (introns) are discarded. For each intron cluster, it iteratively (1) removed introns that were supported with fewer than 100 reads or fewer than 5% of the total number of intronic read counts for the entire cluster, and (2) re-clustered introns according to the procedure above. The intron usage ratio for each clusters was next computed and standardized (across individuals) and quantile normalized (across sample) as in Li et al. ^20^.

#### Association of intron usage with AD and neuropathology traits

The association analysis with neuropathology traits and intron usage was performed using a linear model, adjusting for experimental batch, RNA integrity number (RIN), sex, age at death, and post-mortem interval (PMI). To test for association with AD, we limited the comparison to those participants clinical diagnosis of AD and those who have neither diagnosis (Supplementary Table 1). We used Leafcutter to identify intron clusters with at least one differentially excised intron by jointly modeling intron clusters using a Dirichlet-multinomial GLM^19^. To account for neuronal loss and cell type proportion in each brain sample, we used gene expression level of cell type specific genes as an additional covariate. However, these measures did not affect our association analysis. We report differentially spliced introns at Bonferroni-corrected *P* < 0.05 to correct for multiple hypothesis testing.

We used variancePartition^55^ to estimate the proportion of variance explained of differently excised introns association with AD, burden of amyloid, burden of tangles, and neuritic plaques.

#### Splicing QTL mapping

We used Leafcutter to obtain the proportion of intron defining reads to the total number of reads from the intron cluster it belongs to. This intron ratio describes how often an intron is used relative to other introns in the same cluster. We used WASP^56^ to remove read-mapping biases caused by allele-specific reads. This is particularly significant when a variant is covered by reads that also span intron junctions as it can lead to a spurious association between the variant and intron excision level estimates. We standardized the intron ratio values across individuals for each intron and quantile normalize across introns^57^ and used this as our phenotype matrix. We used linear regression (as implemented in fastQTL)^24^ to test for associations between SNP dosages (MAF ≥ 0.01) within 100kb of intron clusters and the rows of our phenotype matrix that correspond to the intron ratio within each cluster. As covariate, we used the first 3 principal components of the genotype matrix to account for the effect of ancestry plus the first 15 principal components of the phenotype matrix (PSI) to regress out the effect of known and hidden factors. The principal components regress out the technical and biological covariates such as experimental batch, RNA integrity number (RIN), sex, age at death, and post-mortem interval (PMI). To estimate the number of sQTLs at any given false discovery rate (FDR), we ran an adaptive permutation scheme^24^, which maintains a reasonable computational load by tailoring the number of permutations to the significance of the association. We computed the empirical gene-level p-value for the most significant QTL for each gene. Finally, we applied Benjamini-Hochberg correction on the permutation p-values to extract all significant splicing QTL pairs with an FDR < 0.05.

#### Transcriptome-wide Association Studies

We used RNA-seq data and genotypes from ROSMAP to impute the *cis* genetic component of expression/intron usage^36^,^40^ into large-scale late-onset AD GWAS of 74,046 individuals from the International Genomics of Alzheimer’s Project (IGAP)^37^. The complete TWAS pipeline is implemented in FUSION (http://gusevlab.org/projects/fusion/) suite of tools^36^,^40^. The details steps implemented in FUSION are: (1) estimate heritability of gene expression or intron usage unit and stop if not significant. We estimated using a robust version of GCTA-GREML^58^, which generates heritability estimates per feature as well as the as well as the likelihood ratio test (LRT) *P*-value. Only features that have a heritability of Bonferroni-corrected *P* < 0.05 were retained for TWAS analysis. (2) The expression or intron usage weights were computed by modeling all *cis*-SNPs (1MB +/-from TSS) using best linear unbiased prediction (BLUP), or modeling SNPs and effect sizes (BSLMM), LASSO, Elastic Net and top SNPs^36^,^40^. A cross-validation for each of the desired models are performed; (3) Perform a final estimate of weights for each of the desired models and store results. The imputed unit is treated as a linear model of genotypes with weights based on the correlation between SNPs and expression in the training data while accounting for LD among SNPs. To account for multiple hypotheses, we applied an FDR < 0.05 within each expression and splicing reference panel that was used.

We used the same TWAS pipeline to process the CMC datasets (see **Supplementary Notes**).

#### Joint and conditional analysis

Joint and conditional analysis of TWAS results was performed using the summary statistic-based method described in Yang et al.^41^, which we applied to genes instead of SNPs. We used TWAS statistics from the main results and a correlation matrix to evaluate the joint/conditional model. The correlation matrix was estimated by predicting the *cis*-genetic component of expression for each TWAS gene and computing Pearson correlations across all pairs of genes. We used FUSION tool to perform the joint/conditional analysis, generate conditional outputs, and generate plots.

#### Gene Expression, DNA Methylation, Histone Modification QTL Mapping

The details of ROSMAP gene expression, DNA methylation, and histone modification data are described in Supplementary Notes. The quantitative trait locus (xQTL) analysis on a multi-omic dataset is described in Ng et al.^27^. The xQTL results and analysis scripts can be accessed through online portal, xQTL Serve, at http://mostafavilab.stat.ubc.ca/xQTLServe.

#### QTL Sharing

We used the Storey’s Π_1_ statistics^59^ also described in Nica et al.^60^, QTL sharing was estimated as the proportion of true associations Π_1_ among the top SNP in each QTLs in the second QTL.

#### Enrichment of sQTLs within epigenomic marks and splicing factor binding sites

We selected a set of 71 human curated RNA-binding proteins (RBP) splicing regulatory proteins from the SpliceAid-F database^61^ to analyze the relationship between gene expression levels of RBP and intron usage patterns across all samples. To test for enrichment of sQTLs in RBP binding sites, we downloaded human CLIP data in BED format from ClipDB^33^. We used GREGOR^62^ (Genomic Regulatory Elements and Gwas Overlap algoRithm) to evaluate global enrichment of trait-associated variants in splicing factor binding sites. GREGOR^62^ evaluates the significance of the observed overlap (of sQTL and splicing factor binding sites) by estimating the probability of the observed overlap of the lead sQTL relative to expectation using a set of matched control variants (random control SNPs are selected across the genome that match the index SNP for a number of variants in LD, minor allele frequency, and distance to nearest intron).

#### Enrichment of sQTLs in Chromatin States

We downloaded chromatin states from the Roadmap Epigenomics Project. The 15 chromatin states were generated from 5 chromatin marks in DLPFC of a cognitively non-impaired MAP subject with minimal pathology as part of the Roadmap Epigenomics Consortium^25^. A ChromHMM model applicable to brain epigenome was learned by virtually concatenating consolidated data corresponding to the core set of 5 chromatin marks assayed (H3K4me3, H3K4me1, H3K36me3, H3K27me3, H3K9me3). BED files downloaded from http://egg2.wustl.edu/roadmap/web_portal/chr_state_learning.html. To test for enrichment for sQTLs among the 15 chromatin states, we used GREGOR^62^ to evaluate global enrichment of trait-associated variants in splicing factor binding sites.

#### GWAS Enrichment Analyses

We used GARFIELD (unpublished; http://www.ebi.ac.uk/birney-srv/GARFIELD/) to test for enrichment of IGAP AD GWAS SNPs among sQTLs and other publicly available QTL datasets. GARFIELD performs greedy pruning of GWAS SNPs (LD r^2^ >0.1) and then annotates them based on functional information overlap. It quantifies fold enrichment at GWAS *P* <10^−5^ significant cutoff and assesses them by permutation testing, while matching for minor allele frequency, distance to nearest transcription start site and a number of LD proxies (r^2^ > 0.8).

Q-Q plots show quantiles of one dataset against quantiles of a second dataset and are commonly used in GWAS to show a departure from an expected *P*-value distribution. We generated Q-Q plots for LD-pruned GWAS SNPs (PLINK with the settings “--indep-pairwise 100 5 0.8”). We compared the sQTLs overlapping with LD-pruned GWAS SNPs and compared the distribution to a random set of SNPs with similar MAF.

#### GWAS Datasets

We performed transcriptome-wide association study using GWAS summary statistics from: (1) AD GWAS from the International Genomics of Alzheimer’s Project (stage 1 data)^37^; (2) AD genome-wide association study by proxy (GWAx) in 116,196 individuals from the UK Biobank^39^.

#### Protein-protein Interaction Network and Pathway Analysis

We constructed a protein-protein interaction (PPI) network using the GeNets online tool (unpublished; https://apps.broadinstitute.org/genets) to determine whether the AD TWAS genes significantly interact with each other and with known AD associated proteins. GeNets create networks of connected proteins using evidence of physical interaction from the InWeb database, which contains 420,000 high-confidence pair-wise interactions involving 12,793 proteins^63^. Community structures of the underlying genes are displayed in GeNets. These “communities” are also called modules or clusters. This feature highlights genes that are more connected to one another than they are to other genes in other modules. To assess the statistical significance of PPI networks, GeNets applies a within-degree node-label permutation strategy to build random networks that mimic the structure of the original network and evaluates network connectivity parameters on these random networks to generate empirical distributions for comparison to the original network. In addition to PPI network analysis, GeNets allows for gene set enrichment analysis on genes within the PPI network. We used Molecular Signatures Database (MSigDB) Curated Gene Sets (C2), curated from various sources such as online pathway databases, the biomedical literature, and knowledge of domain experts and Canonical Pathways (CP), curated from pathway databases such KEGG, BioCarta, and Reactome to test for gene set enrichment within the PPI network. Then a hypergeometric testing is applied to get *P*-value for gene set enrichment. We used Bonferroni-corrected *P* < 0.05 to correct for multiple hypothesis testing.

#### Data availability

The ROSMAP data are available at the RADC Research Resource Sharing Hub at www.radc.rush.edu. The ROSMAP and MSBB mapped RNA-seq data that support the findings of this study are available in AMP-AD Knowledge Portal (https://www.synapse.org/#!Synapse:syn2580853) upon authentication by the Consortium. The CommonMind Consortium data are available in CMC Knowledge Portal: https://www.synapse.org/#!Synapse:syn4923029.

#### URLs

LeafCutter, https://github.com/davidaknowles/leafcutter;

xQTL Browser, http://mostafavilab.stat.ubc.ca/xQTLServe;

FUSION, http://gusevlab.org/projects/fusion/

MISO, http://genes.mit.edu/burgelab/miso/;

SpliceAid-F, http://srv00.recas.ba.infn.it/SpliceAidF/;

Roadmap Epigenomics Project, http://egg2.wustl.edu/roadmap/web_portal/chr_state_learning.html;

GREGOR, http://genome.sph.umich.edu/wiki/GREGOR;

GARFIELD, http://www.ebi.ac.uk/birney-srv/GARFIELD;

GeNets, https://apps.broadinstitute.org/genets;

Michigan Imputation Server, https://imputationserver.sph.umich.edu/index.html;

The RUSH Alzheimer’s Disease Research Center Research Resource Sharing Hub, https://www.radc.rush.edu;

AMP-AD Synapse Portal, https://www.synapse.org/#!Synapse:syn2580853/wiki/409844;

CommonMind Consortium Knowledge Portal, https://www.synapse.org/#!Synapse:syn2759792/wiki/69613;

IGAP GWAS summary statistics, http://web.pasteur-lille.fr/en/recherche/u744/igap/igap_download.php;

UK Biobank summary statistics, http://gwas-browser.nygenome.org/downloads/gwas-browser/.

## Acknowledgments

We thank the participants of ROS and MAP for their essential contributions and gift to these projects. We thank Alexander Gusev for helpful discussions and for sharing the source code and scripts for TWAS. We thank the International Genomics of Alzheimer’s Project (IGAP) for providing summary results data for these analyses. T.R. is supported by grants from the NIH National Institute on Aging (R01AG054005) and the Alzheimer’s Association. This work has been supported by many different NIH grants: P30AG10161, U01AG046152, R01AG16042, R01AG036836, R01AG015819, R01AG017917, R01AG036547.

We thank the patients and families who donated material for CommonMind Consortium data. The CommonMind Consortium data are available in CMC Knowledge Portal: http://www.synapse.org/#!Synapse:syn4923029. Data were generated as part of the CMC supported by funding from Takeda Pharmaceuticals Company Limited, F. Hoffman-La Roche Ltd and NIH grants R01MH085542, R01MH093725, P50MH066392, P50MH080405, R01MH097276, RO1-MH-075916, P50M096891, P50MH084053S1, R37MH057881 and R37MH057881S1, HHSN271201300031C, AG02219, AG05138 and MH06692. Brain tissue for the study was obtained from the following brain bank collections: the Mount Sinai NIH Brain and Tissue Repository, the University of Pennsylvania Alzheimer’s Disease Core Center, the University of Pittsburgh NeuroBioBank and Brain and Tissue Repositories and the NIMH Human Brain Collection Core. CMC Leadership: Pamela Sklar, Joseph Buxbaum (Icahn School of Medicine at Mount Sinai), Bernie Devlin, David Lewis (University of Pittsburgh), Raquel Gur, Chang-Gyu Hahn (University of Pennsylvania), Keisuke Hirai, Hiroyoshi Toyoshiba (Takeda Pharmaceuticals Company Limited), Enrico Domenici, Laurent Essioux (F. Hoffman-La Roche Ltd), Lara Mangravite, Mette Peters (Sage Bionetworks), Thomas Lehner, Barbara Lipska (NIMH).

## Author Contributions

T.R. and P.L.D. conceived the project and planned the experiments. T.R. and Y.L. analyzed and interpreted the data with support from G.W., S.R., Y.W., I.G. B.N., and S.M. PL.D, D.A.B., M.W., P.S., E.S., V.H., and B.Z. contributed samples and/or data. T.Y.P. performed the *Tau* overexpression in iPSc Neurons. T.R. and P.L.D prepared the first draft of the manuscript. All authors contributed to the final manuscript.

## Corresponding authors

Correspondence to: Towfique Raj and Philip De Jager.

## Reference

1. Wang, E.T. et al. Alternative isoform regulation in human tissue transcriptomes. Nature 456, 470–6 (2008).

2. Kornblihtt, A.R. et al. Alternative splicing: a pivotal step between eukaryotic transcription and translation. Nat Rev Mol Cell Biol 14, 153–65 (2013).

3. Barbosa-Morais, N.L. et al. The evolutionary landscape of alternative splicing in vertebrate species. Science 338, 1587–93 (2012).

4. Wang, G.S. & Cooper, T.A. Splicing in disease: disruption of the splicing code and the decoding machinery. Nat Rev Genet 8, 749–61 (2007).

5. Dredge, B.K., Polydorides, A.D. & Darnell, R.B. The splice of life: alternative splicing and neurological disease. Nat Rev Neurosci 2, 43–50 (2001).

6. Parikshak, N.N. et al. Genome-wide changes in lncRNA, splicing, and regional gene expression patterns in autism. Nature 540, 423–427 (2016).

7. Arai, T. et al. TDP-43 is a component of ubiquitin-positive tau-negative inclusions in frontotemporal lobar degeneration and amyotrophic lateral sclerosis. Biochem Biophys Res Commun 351, 602–11 (2006).

8. Trabzuni, D. et al. MAPT expression and splicing is differentially regulated by brain region: relation to genotype and implication for tauopathies. Hum Mol Genet 21, 4094–103 (2012).

9. Rockenstein, E.M. et al. Levels and alternative splicing of amyloid beta protein precursor (APP) transcripts in brains of APP transgenic mice and humans with Alzheimer's disease. J Biol Chem 270, 28257–67 (1995).

10. Buee, L., Bussiere, T., Buee-Scherrer, V., Delacourte, A. & Hof, P.R. Tau protein isoforms, phosphorylation and role in neurodegenerative disorders. Brain Res Brain Res Rev 33, 95–130 (2000).

11. Valenca, G.T. et al. The Role of MAPT Haplotype H2 and Isoform 1N/4R in Parkinsonism of Older Adults. PLoS One 11, e0157452 (2016).

12. Bai, B. et al. U1 small nuclear ribonucleoprotein complex and RNA splicing alterations in Alzheimer's disease. Proc Natl Acad Sci U S A 110, 16562–7 (2013).

13. Vaquero-Garcia, J. et al. A new view of transcriptome complexity and regulation through the lens of local splicing variations. Elife 5, e11752 (2016).

14. Lambert, W. et al. Probing the transient interaction between the small heat-shock protein Hsp21 and a model substrate protein using crosslinking mass spectrometry. Cell Stress Chaperones 18, 75–85 (2013).

15. Raj, T. et al. Polarization of the effects of autoimmune and neurodegenerative risk alleles in leukocytes. Science 344, 519–23 (2014).

16. Raj, T. et al. CD33: increased inclusion of exon 2 implicates the Ig V-set domain in Alzheimer's disease susceptibility. Hum Mol Genet 23, 2729–36 (2014).

17. Bennett, D.A., Schneider, J.A., Arvanitakis, Z. & Wilson, R.S. Overview and findings from the religious orders study. Curr Alzheimer Res 9, 628–45 (2012).

18. Bennett, D.A. et al. Selected findings from the Religious Orders Study and Rush Memory and Aging Project. J Alzheimers Dis 33 Suppl 1, S397–403 (2013).

19. Li, Y.I., Knowles, D.A. & Pritchard, J.K. LeafCutter: Annotation-free quantication of RNA splicing. bioArxiv (2017).

20. Li, Y.I. et al. RNA splicing is a primary link between genetic variation and disease. Science 352, 600–4 (2016).

21. Tollervey, J.R. et al. Analysis of alternative splicing associated with aging and neurodegeneration in the human brain. Genome Res 21, 1572–82 (2011).

22. Mitchelmore, C. et al. NDRG2: a novel Alzheimer's disease associated protein. Neurobiol Dis 16, 48–58 (2004).

23. AMP-AD Knowledge Portal. https://whttp://www.synapse.org/ -!Synapse:syn2580853/wiki/409840.

24. Ongen, H., Buil, A., Brown, A.A., Dermitzakis, E.T. & Delaneau, O. Fast and efficient QTL mapper for thousands of molecular phenotypes. Bioinformatics 32, 1479–85 (2016).

25. Bernstein, B.E. et al. The NIH Roadmap Epigenomics Mapping Consortium. Nat Biotechnol 28, 1045–8 (2010).

26. Fromer, M. et al. Gene expression elucidates functional impact of polygenic risk for schizophrenia. Nat Neurosci 19, 1442–1453 (2016).

27. Ng, B. et al. Brain xQTL map: integrating the genetic architecture of the human brain transcriptome and epigenome bioArxiv (2017).

28. Nicolae, D.L. et al. Trait-associated SNPs are more likely to be eQTLs: annotation to enhance discovery from GWAS. PLoS Genet 6, e1000888 (2010).

29. Chen, L. et al. Genetic Drivers of Epigenetic and Transcriptional Variation in Human Immune Cells. Cell 167, 1398–1414 e24 (2016).

30. Malik, M. et al. CD33 Alzheimer's risk-altering polymorphism, CD33 expression, and exon 2 splicing. J Neurosci 33, 13320–5 (2013).

31. Huang, K.L., Marcora, E., Pimenova, A. & et al. A common haplotype lowers SPI1 (PU.1) expression in and delays age at onset for Alzheimer's disease. Nat. Neuroscience, In Press. (2017).

32. Sibley, C.R., Blazquez, L. & Ule, J. Lessons from non-canonical splicing. Nat Rev Genet 17, 407–21 (2016).

33. Yang, Y.C. et al. CLIPdb: a CLIP-seq database for protein-RNA interactions. BMC Genomics 16, 51 (2015).

34. Scheckel, C. et al. Regulatory consequences of neuronal ELAV-like protein binding to coding and non-coding RNAs in human brain. Elife 5(2016).

35. Borreca, A., Gironi, K., Amadoro, G. & Ammassari-Teule, M. Opposite Dysregulation of Fragile-X Mental Retardation Protein and Heteronuclear Ribonucleoprotein C Protein Associates with Enhanced APP Translation in Alzheimer Disease. Mol Neurobiol 53, 3227–3234 (2016).

36. Gusev, A. et al. Integrative approaches for large-scale transcriptome-wide association studies. Nat Genet 48, 245–52 (2016).

37. Lambert, J.C. et al. Meta-analysis of 74,046 individuals identifies 11 new susceptibility loci for Alzheimer's disease. Nat Genet 45, 1452–8 (2013).

38. Vance, C. et al. Mutations in FUS, an RNA processing protein, cause familial amyotrophic lateral sclerosis type 6. Science 323, 1208–11 (2009).

39. Liu, J.Z., Erlich, Y. & Pickrell, J.K. Case-control association mapping by proxy using family history of disease. Nat Genet 49, 325–331 (2017).

40. Gusev, A., et al. Transcriptome-wide association study of schizophrenia and chromatin activity yields mechanistic disease insights. bioArxiv.

41. Yang, J. et al. Conditional and joint multiple-SNP analysis of GWAS summary statistics identifies additional variants influencing complex traits. Nat Genet 44, 369-75, S1-3 (2012).

42. Raj, T. et al. Alzheimer disease susceptibility loci: evidence for a protein network under natural selection. Am J Hum Genet 90, 720–6 (2012).

43. Gotzl, J.K., Lang, C.M., Haass, C. & Capell, A. Impaired protein degradation in FTLD and related disorders. Ageing Res Rev 32, 122–139 (2016).

44. Nixon, R.A. New perspectives on lysosomes in ageing and neurodegenerative disease. Ageing Res Rev 32, 1 (2016).

45. Emmett, M.J. et al. Histone deacetylase 3 prepares brown adipose tissue for acute thermogenic challenge. Nature 546, 544–548 (2017).

46. Tian, Y., Chang, J.C., Fan, E.Y., Flajolet, M. & Greengard, P. Adaptor complex AP2/PICALM, through interaction with LC3, targets Alzheimer's APP-CTF for terminal degradation via autophagy. Proc Natl Acad Sci U S A 110, 17071–6 (2013).

47. Bennett, D.A. et al. Overview and findings from the rush Memory and Aging Project. Curr Alzheimer Res 9, 646–63 (2012).

48. Purcell, S. et al. PLINK: a tool set for whole-genome association and population-based linkage analyses. Am J Hum Genet 81, 559–75 (2007).

49. Patterson, N., Price, A.L. & Reich, D. Population structure and eigenanalysis. PLoS Genet 2, e190 (2006).

50. Das, S. et al. Next-generation genotype imputation service and methods. Nat Genet 48, 1284–7 (2016).

51. McCarthy, S. et al. A reference panel of 64,976 haplotypes for genotype imputation. Nat Genet 48, 1279–83 (2016).

52. Dobin, A. et al. STAR: ultrafast universal RNA-seq aligner. Bioinformatics 29, 15–21 (2013).

53. Li, B. & Dewey, C.N. RSEM: accurate transcript quantification from RNA-Seq data with or without a reference genome. BMC Bioinformatics 12, 323 (2011).

54. Johnson, W.E., Li, C. & Rabinovic, A. Adjusting batch effects in microarray expression data using empirical Bayes methods. Biostatistics 8, 118–27 (2007).

55. Hoffman, G.E. & Schadt, E.E. variancePartition: interpreting drivers of variation in complex gene expression studies. BMC Bioinformatics 17, 483 (2016).

56. van de Geijn, B., McVicker, G., Gilad, Y. & Pritchard, J.K. WASP: allele-specific software for robust molecular quantitative trait locus discovery. Nat Methods 12, 1061–3 (2015).

57. Degner, J.F. et al. DNase I sensitivity QTLs are a major determinant of human expression variation. Nature 482, 390–4 (2012).

58. Yang, J., Lee, S.H., Goddard, M.E. & Visscher, P.M. GCTA: a tool for genome-wide complex trait analysis. Am J Hum Genet 88, 76–82 (2011).

59. Storey, J.D. & Tibshirani, R. Statistical significance for genomewide studies. Proc Natl Acad Sci U S A 100, 9440–5 (2003).

60. Nica, A.C. et al. The architecture of gene regulatory variation across multiple human tissues: the MuTHER study. PLoS Genet 7, e1002003 (2011).

61. Giulietti, M. et al. SpliceAid-F: a database of human splicing factors and their RNA-binding sites. Nucleic Acids Res 41, D125–31 (2013).

62. Schmidt, E.M. et al. GREGOR: evaluating global enrichment of trait-associated variants in epigenomic features using a systematic, data-driven approach. Bioinformatics 31, 2601–6 (2015).

63. Lage, K. et al. A human phenome-interactome network of protein complexes implicated in genetic disorders. Nat Biotechnol 25, 309–16 (2007).

64. Fairfax, B.P. et al. Innate immune activity conditions the effect of regulatory variants upon monocyte gene expression. Science 343, 1246949 (2014).

